# A lateralised design for the interaction of visual memories and heading representations in navigating ants

**DOI:** 10.1101/2020.08.13.249193

**Authors:** Antoine Wystrach, Florent Le Moël, Leo Clement, Sebastian Schwarz

## Abstract

The navigational skills of ants, bees and wasps represent one of the most baffling examples of the powers of minuscule brains. Insects store long-term memories of the visual scenes they experience ^1^, and they use compass cues to build a robust representation of directions ^2,3^. We know reasonably well how long-term memories are formed, in a brain area called the Mushroom Bodies (MB) ^4–8^, as well as how heading representations are formed in another brain area called the Central Complex (CX) ^9–12^. However, how such memories and heading representations interact to produce powerful navigational behaviours remains unclear ^7,13,14^. Here we combine behavioural experiments with computational modelling that is strictly based on connectomic data to provide a new perspective on how navigation might be orchestrated in these insects. Our results reveal a lateralised design, where signals about whether to turn left or right are segregated in the left and right hemispheres, respectively. Furthermore, we show that guidance is a two-stage process: the recognition of visual memories – presumably in the MBs – does not directly drive the motor command, but instead updates a “desired heading” – presumably in the CX – which in turn is used to control guidance using celestial compass information. Overall, this circuit enables ants to recognise views independently of their body orientation, and combines terrestrial and celestial cues in a way that produces exceptionally robust navigation.

## Bilaterally decorrelated input to the CX produces goal-oriented paths

We first investigated how information about the visual familiarity of the scenes – as computed in the MB – could plausibly be sent to the CX for guidance given the known circuitry of insect brains. Even though our current approach is an experimental one, the CX circuitry is understood and conserved enough to make such effort possible using biologically constrained neural modelling ^12,15–17^.

Recent studies have shown that the CX circuits can: 1) track the current heading, in two substructures called Ellipsoid Body (EB) and Protocerebral Bridge (PB) ^10,11,18^; 2) retain a desired heading representation for tens of seconds in the Fan-shaped Body (FB) ^14^; and 3) compare both current and desired headings to output compensatory left/right steering commands ^14,19^. The desired heading can be updated by bilateral signals to the FB from external regions ^14^. Such a signal can plausibly come from the recognition of long-term visual memories in the MB, which sends bilateral input to the FB through one relay in the Superior Intermediate Protocerebrum (SIP). These observations led to the idea that navigation, such as learnt route following, could emerge by having the MBs signalling to the CX when the insect is facing its familiar route direction or not ^7,13,14^.

We thus tested the viability of this hypothesis by building a model of the CX, strictly based on this connectivity (Fig. 1). Contrary to what was expected, our model shows that having the bilateral ‘visual familiarity’ signals to the FB correspond with the moments when the agent is facing the correct route direction did not allow straight routes to emerge. A thorough search through the parameter space revealed that this configuration produces a mediocre directionality at best, and is very sensitive to parameter change (Extended data fig. 1). Contrastingly, route following becomes extremely stable and robust to parameter changes as soon as the signals to the FB from the left and right brain hemispheres correspond to moments where the agent is oriented to the right or the left of its goal, respectively (Fig. 1). Impressively, varying parameters (such as the time during which FB neurons sustain their activity, or the heading angle away from the goal for which left or right input signals are strongest) hardly has any effect: straight routes emerge as long as left and right hemispheric inputs roughly correlate with a right and left heading bias, respectively (Fig. 1, Extended data fig. 1). As a corollary, if left and right hemispheric inputs correlate instead with left and right (rather than right and left) heading biases, a straight route in the reverse direction emerges (Fig. 1). Thus, having the input signal correlate with moments where the agent faces the goal direction corresponds to a zone of transition between two stable regimes of route-following in opposite directions.

**Figure 1.**
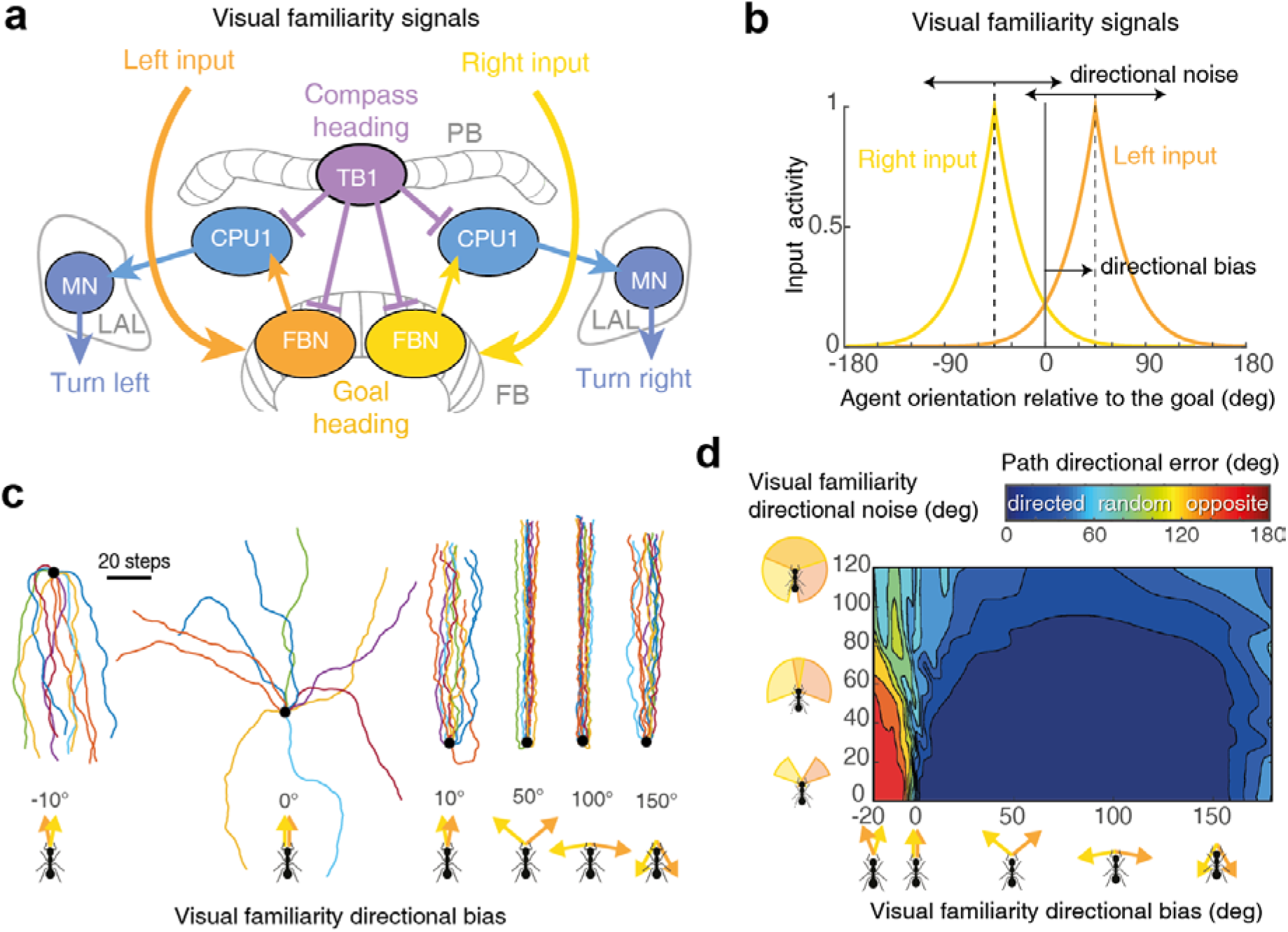
Bilaterally decorrelated input to Central Complex produces stable route heading. ***a***. *The* central complex (CX) sits at the centre of the brain but is wired to both hemispheres. It receives bilateral inputs in the Fan-shaped Body (FB), where sustained activity of the FB neurons (FBN) forms two representations of the goal heading. CPU1 neurons compare such ‘goal heading’ representations to the ‘compass-based current heading’ representation of the Protocerebral Bridge (PB) neurons (TB1) and outputs bilateral signals to the left and right Lateral Accessory Lobes (LALs), where they modulate motor neurons (MN) descending to the thorax to control left and right turns, respectively (see **extended figure 2, d, g** for details of the circuitry). **b**. Simulated inputs to the FBN neurons. We assumed that the input signals to the FBN are body-orientation-dependant (as expected if resulting from visual familiarity of the scene ^28^ such as outputted by the MBs ^4^. ‘directional bias’ indicates the direction relative to the goal direction (0°) at which the left visual familiarity signals is highest in average (+45° in this example). Right signal responds symmetrically for the other direction (-directional bias). ‘Directional noise’ in the visual familiarity was implemented by shifting the input curve response around its mean (i.e. the ‘directional bias’) at each time step by a random value (normal distribution with standard deviation given by ‘directional noise’). **c**. Paths resulting given different directional biases. **d**. Path directional error (absolute angular error between start-to-arrival beeline, and start-to-goal direction) after 200 steps, as a function of the visual familiarity ‘directional bias’ (x axis) and ‘directional noise’ (y axis). **c, d**. Straight route headings robustly emerge as long as left and right inputs send a signal when the body is oriented right and left from the goal, respectively (i.e., directional bias > 0°) but not if both inputs send a signal when facing the goal (i.e., directional bias = 0°). Orientation towards the opposite direction emerges if left and right inputs signal inversely, that is, when the body is oriented right and left from the goal respectively (i.e., directional bias < 0°). Robustness to visual familiarity directional noise indicate that the direction in which views are learnt does not need to be precisely controlled. See further analysis in Extended Data Fig. 1.

In other words, this suggests that recognising views when facing the goal may not be a good solution, and instead, it shows that the CX circuitry is remarkably adapted to control a visual course as long as the input signals from the visual familiarity of the scene to both hemispheres are distinct, with one hemisphere signalling when the agent’s heading is biased towards the right and the other, towards the left. This model makes particular predictions, which we next tested with behavioural experiments.

## The recognition of familiar views triggers compensatory left or right turns

Previous studies assumed that ants memorise views while facing the goal ^20–22^ and anti-goal ^23–25^) directions, and that they must consequently align their body in these same directions to recognise a learnt view as familiar ^26–28^. On the contrary, our modelling effort suggests that ants should rather recognise views based on whether the route direction stands on their ‘left or right’ rather than ‘in front or behind’. We put this idea to the test using an open-loop trackball system enabling the experimenter to choose both the position and body orientation of tethered ants directly in their natural environment ^29^. We trained ants along a route and captured homing individuals just before they entered their nest to ensure that these so-called zero-vector ants (ZV) could no longer rely on their path integration homing vector ^30^. We recorded the motor response of these ants while mounted on the trackball system, in the middle of their familiar route, far from the catchment area of the nest, when fixed in eight different body orientations (Fig. 2a, b). Results show that, irrespective of their body orientation, ants turned mostly towards the correct route direction (Fig. 2c). When the body was oriented towards (0°, nest direction) or away (180°) from the route direction, ants still showed a strong preference for turning on one side (to the left or to the right, depending on individuals) (Fig. 2d). This was not the case when ants were tested in unfamiliar surroundings (Fig. 2c, d), showing that the lateralised responses observed on the familiar route was triggered by the recognition of the visual scene. This implies that ants can recognise their route independently of their body orientation, and can derive whether the route direction is towards their left or their right. Importantly, even when facing the route or anti-route direction, recognition of familiar views appears to trigger a ‘left vs. right’ decision rather than a ‘go forward vs. turn’ decision.

**Figure 2.**
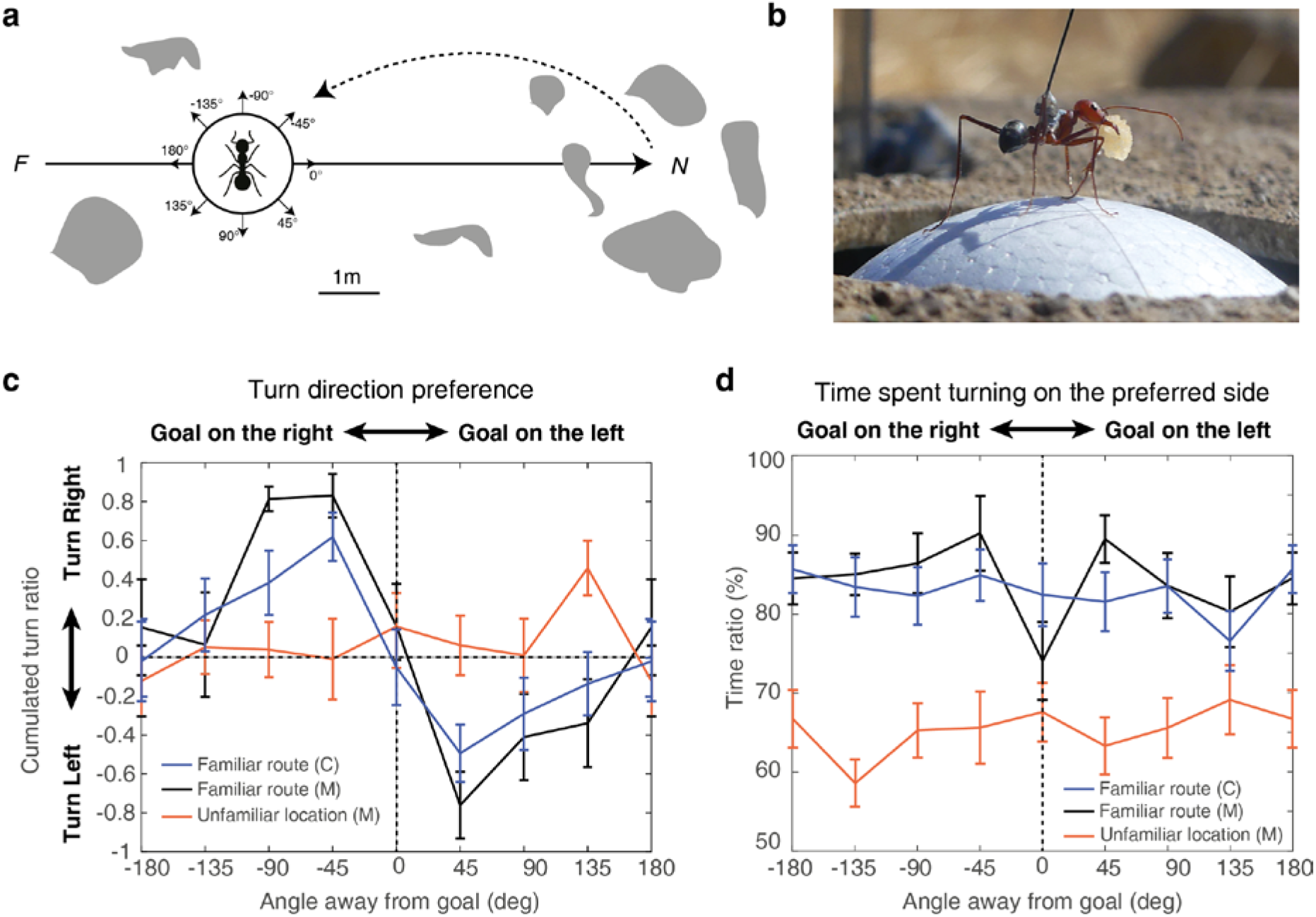
Ants visually recognise whether the goal direction is left or right. **a**. Homing ants were captured at the end of their familiar route and fixed on the trackball (**b**) in 8 different compass orientations. The route was rich in visual terrestrial cues (grey blobs). *F*: feeder, *N*: nest. **b**. An individual *Cataglyphis velox* mounted on the trackball setup, holding its precious cookie crumb. **c**. Turn ratio (degrees *(right − left) / (right + left)*; mean ± se across individuals) for the eight compass directions, on the familiar route or in the unfamiliar location (same compass directions but unfamiliar surroundings) across 12 seconds of recording. **d**. Proportion of time spent turning on the preferred side of each individual (mean ± se across individuals). C: *Cataglyphis velox (n=17)*, M: *Myrmecia crosslandi (n=11*).

## Guidance based on memorised views involves the celestial compass

We showed that the recognition of familiar views indicates whether the goal direction is towards the left or right. In principle, guidance could thus be achieved by having these left/right signals directly trigger the left or right motor command. An alternative would be, as in our model, that such left/right signals can be used to update the ‘desired heading directions’ in the CX, which in turn uses its own compass information to control steering (Fig. 1). This makes a counterintuitive prediction: if the recognition of familiar views triggers a turn towards the correct side, reversing the direction of the compass representation in the CX should immediately reverse the motor decision. We tested this prediction by mirroring the apparent position of the sun in the sky by 180° to *Cataglyphis velox* ants tethered to our trackball system. A previous study had shown that this manipulation was sufficient to shift this species’ compass heading representation ^31^.

We first tethered well-trained ZV ants (i.e., captured just before entering the nest) on our trackball system with their body orientation fixed perpendicularly to their familiar route direction. As expected, ants in this situation turned towards the correct route direction (Fig. 3, left panels, natural sun), indicating that they correctly recognised familiar visual terrestrial cues. When mirroring the apparent sun’s position by 180°, these ants responded by turning in the opposite direction within one second (Fig. 3, left panel, mirrored sun). We repeated the experiment by placing such ZV ants in the same compass direction but in an unfamiliar location. In this situation, the ants turned in random directions (Fig. 3, middle panels), showing that the direction initially chosen by the ants on their familiar route (Fig. 3, left panels) was based on the recognition of terrestrial rather than celestial cues. It however remains unclear whether the sun rotation had an impact on ants in unfamiliar terrain, as ants in this situation regularly alternate between left and right turns anyway ^25^. Finally, to ensure that the observed effect on route was not due to an innate bias at this particular location, we repeated this experiment with ants tethered at the exact same route location and body orientation, but this time only with ants that were trained to an alternative straight route, which was aligned with the tethered direction of the trackball (Fig. 3, right panel). As expected, these ants showed no preference in turning direction at the group level, although most individuals still strongly favoured one side rather than walking straight (Fig. 3 right panels). Interestingly, mirroring the sun significantly reversed the individual’s chosen direction (even though they were aligned with their goal direction) (Fig. 3c right panels).

**Figure 3.**
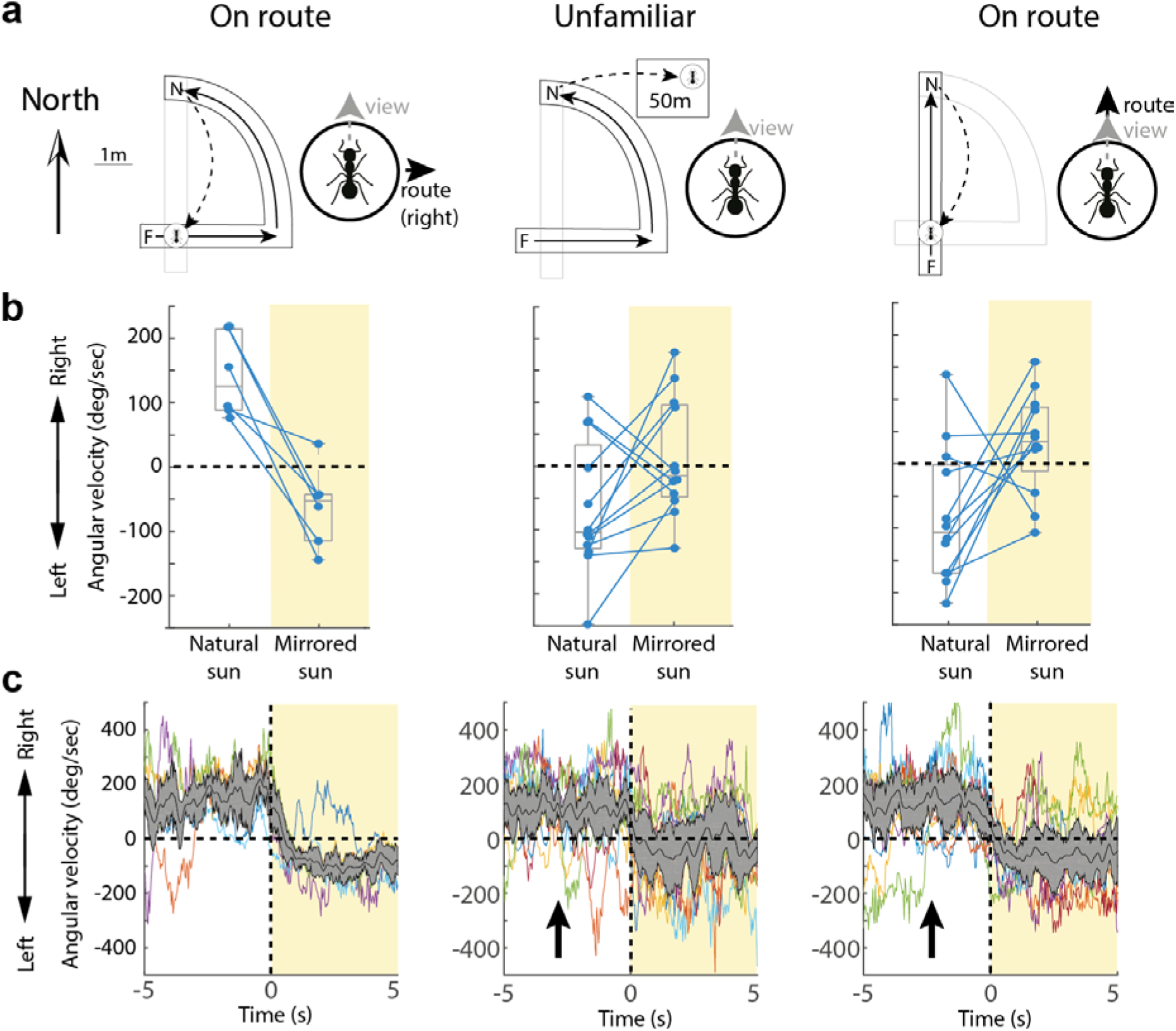
Rotation of celestial cues shift turning direction based on familiar terrestrial cues. **a**. Schemes of the training and test condition. Homing ants were captured at the end of their familiar route (black arrows: familiar route, F: feeder, N: nest) and fixed on the trackball with their body always facing north, either on their route with the route direction 90° to the right (left panel); or within unfamiliar surroundings (middle panel); or ants were trained along a route oriented 90° to the previous one and released on their familiar route in the same location and orientation, which this time is facing their route direction. **b**. Box plots indicate average angular velocity (positive = right turn) each ant (dots) 5s before (white) and 5s after (yellow) the apparent sun’s position is mirrored by 180°. Wilcoxon test for: ‘turn towards the right with natural sun’ (left panel: n=6, p=0.0156; middle panel: n=12 p=0.9788; right panel: n=12 p=0.9866), ‘mirror effect: turn direction reversal’ (left panel: n=6, p=0.0156, power=0.9994; middle panel: n=12 p=0.3955; right panel: n=12 p=0.0320). **c**. Turning velocities (individuals in colour; median ± iqr of the distribution in grey) across time, before and after the sun manipulation (t_0_). Arrows in the middle and left panels: the velocities of some individuals have been inverted so that all individuals’ mean turn directions before the manipulation are positive.

Taken together these results show that guidance based on learnt views is a two-stage process: the recognition of visual memories – presumably through the MBs – does not directly drive the motor command, but it instead signals a desired heading – presumably through the CX –, which in turn is used to control guidance using celestial compass information.

### A complex interaction between terrestrial and celestial guidance

The results from above point at a complex interaction between the use of long-term memory of terrestrial cues – indicating whether the goal is left or right – and the heading estimate based on compass cues. To further endorse the credibility of our proposed guidance system, we used our model to explore how agents navigating along their familiar route would react to a sudden 135° shift of the CX current celestial compass estimate, and compared their behaviour to that of real homing ZV ants tested in a similar scenario, where we shifted the sun position by 135° using a mirror (Fig. 4). Impressively, and despite the nonlinear dynamics at play, the simulated shift in the CX model closely resembled the response of the ants to the sun manipulation, adding credibility to the model and helping us grasp the mechanisms at play (Fig. 4).

**Figure 4.**
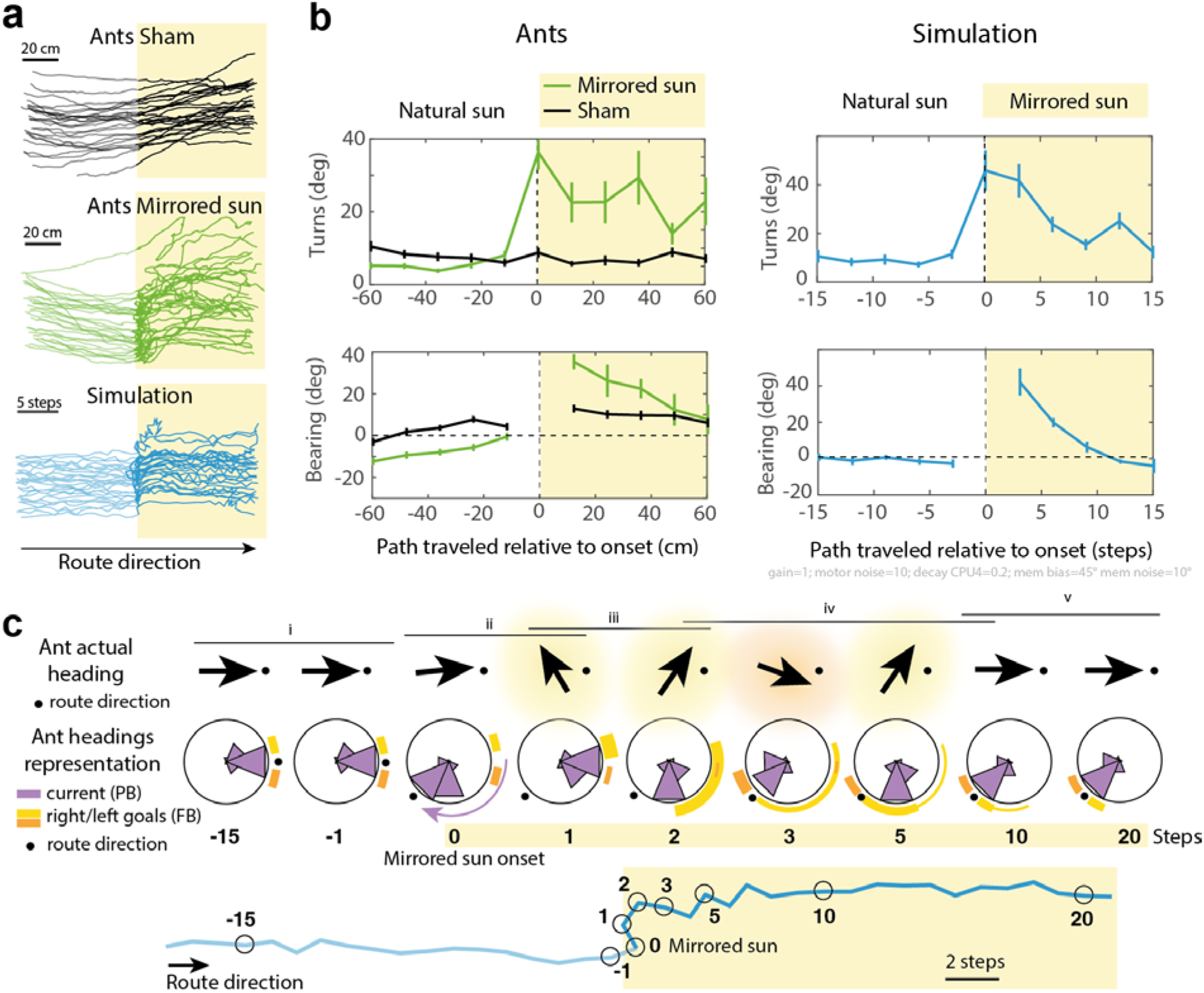
Rotation of celestial cues affect ants route following as predicted. Paths (**a**) and quantification of bearing and turns (**b**) of real (black and green) and simulated (blue) zero-vector ants (i.e., deprived of path integration information) recapitulating a familiar straight route while entering an area where we manipulated celestial compass cues (yellow). For the ‘Mirrored sun’ condition (green) the real sun was hidden from the ants and mirrored so as to appear rotated by 135° counter clockwise in the sky. For the ‘sham’ condition (black), the experimenters were standing in the same place and the real sun was also hidden, but only a small piece of the sky (close to, but not including the sun) was mirrored for the ants. Simulated ants (blue) result from the model presented in fig. 1. Sun rotation was modelled as a 135° shift in the current heading representation (3-cell shift of the bump of activity in the Protocerebral Bridge). Paths of both real and simulated ants were discretised (segments of 12 cm for real ants, and of 3 steps for the simulations), before and after the sun rotation onset point. Turns correspond to the absolute angle between two successive segments, bearing indicates the direction of segments relative to the route (0°). Turns at ‘0’ on the x-axes correspond to the angle between the segment preceding and following the shift of the celestial compass. **c**. The effect observed in the simulations is quantitatively dependant on the model’s parameters (here gain=1; motor noise=10; decay FBN=0.2; visual familiarity directional bias ± noise=45°±10° see Extended Data Fig. 1 for a description of parameters), but its key signature can be explained qualitatively. (i) Under normal situation the current heading is maintained between the right and left goal heading representation in the Fan-shaped Body (FB) (yellow and orange marks) and updated by right and left visual familiarity signals. (ii) The sun rotation creates a sudden shift of the current heading representation in the Protocerebral Bridge (PB) (purple curved arrow), although the agent is still physically facing the actual route direction (black dot). This leads the agent to display a sudden left turn to re-align its shifted heading representation with the FB goal heading that is held in short term memory. (iii) This novel direction of travel is visually recognised as being ‘left of the goal’, causing a strong lateralised signal in the right FB’s goal heading representation (yellow). This biased activity triggers right turns, exposing the agent to new headings recognised as ‘right of the goal’, and thus more signal sent to the right FB (yellow arcs), favouring further right turns. (iv) Turning right eventually leads the agent to overshoot the actual goal direction, recognise view as ‘right from the goal’ and thus signalling in the left FB (orange). These signals are, at first, superimposed with the previous desired heading representation, resulting in a period of conflicting guidance information causing meandering. (v) The agent progressively updates its novel goal heading representation as the trace of the previous desired heading fades out and the new one strengthens due to the incoming signals from visual familiarity. In sum, motor decision results from complex dynamics between two main factors: 1- how strong are the left and right visual familiarity signals updating the goal heading representations (orange and yellow glow around the ‘Ant actual heading’ arrows), which depend on whether the agent is oriented left or right from its goal; and 2- how well the current heading representation (PB) matches the goal heading representation (more detail in Extended Data Fig. 2).

### General discussion

We showed that during view-based navigation, ants recognise views when oriented left and right from their goal to trigger left and right turns. Facing in the correct route direction does not trigger a ‘go forward’ command, but marks some kind of labile equilibrium point in the system. Also, we show that the recognition of left or right familiar views does not drive the motor decision directly but is perfectly suited to inform the CX, which in turn maintains the desired heading using its own compass information. The advantage of this design is clear considering that the recognition of learnt visual terrestrial cues is sensitive to variables such as body orientation ^31,32^ or partial visual obstructions that must happen continuously when navigating through grassy or leafy environments, making the visual familiarity signal mediated by the MBs inherently noisy. In contrast, the CX provides a stable and sustained heading representation by integrating self-motion ^11^ with multiple wide-field celestial ^10^ and terrestrial cues ^9,33^. The CX is thus well suited to act as a heading buffer from the noisy MBs signal, resulting in smooth and stable guidance control. In addition, the compass representation in the CX enables to steer the direction of travel independently of the actual body orientation ^12^. Our results thus explain how ants visually recognise a view using the MBs and subsequently follow such direction backwards using the CX ^31^ or how ants can estimate the actual angular error between the current and goal directions before initiating their turn ^34^. Also, in addition to route following, such a lateralised design can produce remarkably robust homing in complex environments (Wystrach et al., 2020 in prep).

Finally, the proposed circuit offers an interesting take on the evolution of navigation. Segregating ‘turn left’ and ‘turn right’ signals between hemispheres evokes the widespread tropotaxis, where orientation along a gradient is achieved by directly comparing the signals intensities between physically distinct left and right sensors (e.g., antennae or eyes) in bilateral animals ^35–41^. Comparing signals between hemispheres could thus be an ancestral strategy in arthropods; and ancestral brain structures such as the CX accommodates well such a bilateral design and may be constrained to receive such lateralised input to function properly. The evolution of visual route-following in hymenoptera is a relatively recent adaptation, and it cannot be achieved by directly comparing left and right visual inputs – which is probably why each eye can afford to project to both hemispheres’ MBs ^42,43^. Categorising learnt views as indicators of whether the goal is to the left or to the right, and subsequently segregating this information in the left and right hemispheres may thus be an evolutionary adaptation to fit the ancestrally needed bilateral inputs to the CX (Fig. 1).

How left and right visual memories are acquired and learnt when naive insects explore the world for the first time remains to be seen. During their learning flights, wasps regularly alternate between moments facing 45° to the left and 45° to the right of their goal, strongly supporting our claim that insect form such left and right memories ^44^. During their meandering learning walks, ants tend to reverse turning direction when facing the nest or anti-nest direction ^21,23,45^, however, they do expose their gaze in all directions, providing ample opportunities to form a rich set of left and right visual memories ^45^. Our model shows that the angle at which views are learnt does not need to be precisely controlled (Fig. 1c,d). Views facing the nest may as well be included during learning and categorised as left, right or both, explaining why most ants facing their goal usually choose to turn in one particular direction while others turned less strongly. During learning, the first source of information about whether the current body orientation is left or right from the goal probably results from path integration. Interestingly, lateralised dopaminergic feedback from the Lateral Accessory Lobes (LAL, a pre-motor area) to the MBs could represent an ideal candidate to orchestrate such a categorisation of left/right memories (Wystrach et al., 2020 in prep). Revisiting current questions in insect and robot navigation such as early exploration, route following and homing ^20,46–49^; the integration of aversive memories ^8,24,50^, path integration and views (^51–54^ or other sensory modalities (^55–58^ as well as seeking for underlying neural correlates ^5–7^ – with such a lateralised design as a framework promises an interesting research agenda.

## Acknowledgments

We thank the Profs. Jochen Zeil and Xim Cerda to provide us access to field sites in ANU Canberra, Australia and Sevilla, Spain, respectively. We are grateful to the Prof. Hansjuergen Dahmen for helping us setting the trackball device. We thank the Profs. Rüdiger Wehner, Tom Collett and Paul Graham for fruitful discussions and comments on earlier versions of the manuscript.

## Authors contributions

Research design: AW. Data collection: AW, SS, FLM, LC. Trackball system design: FLM. Data analysis: AW, SS, FLM. Modelling: AW. Manuscript writing: AW.

## Method

### The trackball setup

For both experiments (fig 2 and 3) we used the air-suspended trackball setup as described in Dahmen et al., 2017 ^29^; and chose the configuration where the ants are fixed in a given direction and cannot physically rotate (if the ant tries to turn, the ball counter-rotates under its legs). To fix ants on the ball, we used a micro-magnet and metallic paint applied directly on the ant’s thorax. The trackball air pump, battery and computer were connected to the trackball through 10 m long cables and hidden in a remote part of the panorama. The trackball movements were recorded using custom software in C++, data was analysed with Matlab and can be provided upon request.

### Routes setups and ant training in Cataglyphis velox

For all experiments (fig. 2 and 3 and 4), *Cataglyphis velox* ants were constrained to forage within a route using dug wood planks that prevented them to escape, while leaving the surrounding panoramic view of the scenery intact (as described in Wystrach et al., 2012^59^). Cookie crumbs were provided ad libitum in the feeder positions for at least two days before any tests. Some barriers dug into the ground created baffles, enabling us to control whether ants were experienced with the route. Ants were considered trained when able to home along the route without bumping into any such obstacle. These ants were captured just before they entered their nest to ensure that they could not rely on path integration (so-called ZV ants), marked with a metallic paint on the thorax and a colour code for individual identification, and subjected to tests (see next sections).

### Routes setups and ant training in Myrmecia croslandi

For the experiment with *Myremcia croslandi* ants (fig. 2), we used each individual’s natural route, for which these long-lived ants have extensive experience ^60^. Individuals were captured on their foraging trees, marked with both metallic paint and a colour code for individual identification, given a sucrose solution or a prey and released where they had been captured (on their foraging tree). Upon release, most of these ants immediately started to return home. We followed them while marking their route using flag pins every 50 cm (so that their exact route was known). We captured the ants just before they entered their nests and subjected them to the test on the trackball (see next section).

### Experimental protocol for the left/right trackball experiment (figure 2)

1- An experienced ant was captured just before entering its nest, and marked with a drop of metallic paint on the thorax.

2- A large opaque ring (30 cm diameter, 30 cm high) was set around the trackball setup.

3- The ant was fixed on the trackball within the opaque ring, which prevented her to see the surroundings. Only a portion of sky above was accessible to the ant.

4- The trackball system (together with the opaque ring and the fixed ant within) was moved to the desired position and rotated so that the ant was facing the desired direction.

5- One experimenter started recording the trackball movements (from the remote computer), when another lifted the ring (so the ant could see the scenery) before leaving the scene, letting the ant behave for at least 15 seconds post ring lifting.

6- The experimenter came back, replaced the ring around the trackball system, and rotated the trackball system (following a pre-established pseudo random sequence) for the ant to face in a novel direction.

7- We repeated steps 5 and 6 until the 8 possible orientations were achieved (the sequence of orientations were chosen in a pseudo-random order so as to counter-balance orientation and direction of rotation).

The data shown in fig. 2 for each orientation is averaged across 12 sec of recording (from 3 sec to 15 sec assuming ring lifting is at 0 sec). We decided to let 3 sec after ring lifting, as the movements of the experimenter before he leaves the scenery might disturb the ants).

In all experiments, ants were tested only once.

### Experimental protocol for the mirror trackball experiments (figure 3)

1- An experienced ant was captured just before entering its nest, and marked with a drop of metallic paint on the thorax.

2- A large opaque ring (30 cm diameter, 30 cm high) was set around the trackball setup.

3- The ant was fixed on the trackball within the opaque ring, which prevented her to see the surroundings. Only a portion of sky above was accessible to the ant.

4- The trackball system (together with the opaque ring and the fixed ant within) was moved to the desired position and rotated so that the ant was facing the desired direction.

5- One experimenter started recording the trackball movements, when another lifted the ring (so the ant could see the scenery) before leaving the scene, letting the ant behave for at least 10 seconds post ring lifting.

5- Two experimenters simultaneously hid the real sun and projected the reflected sun using a mirror, so that the sun appeared in the opposite position of the sky to the ant for at least 8 seconds.

Ants were tested only once, in one of the conditions.

### *Experimental design and protocol for the mirror experiment* with ants on the floor *(figure 4)*

*Cataglyphis velox* ants were trained to a 10 meters-long route for at least two consecutive days. A 240 × 120 cm thin wood board was placed on the floor in the middle of the route, ensuring that the navigating ants walked smoothly without encountering small clutter over this portion of the route. Homing ants were captured just before entering their nest and released at the feeder as ZV ants. Upon release, these ZV ants typically resume their route homing behaviour; at mid-parkour (halfway along the board section) the real sun was hidden by one experimenter and reflected by another, using a mirror, for the sun to appear to the ant 135° away from its original position in the sky. To ensure that each individual was tested only once, tested ants were marked with a drop of paint after the procedure.

The ZV ants walking on the board were recorded using a Panasonic Lumix DMC-FZ200 camera on a tripod, and their paths were digitised frame by frame at 10 fps using image J. We used four marks on the board to correct for the distortion due to the tilted perspective of the camera’s visual field. Analysis of the paths were achieved with Matlab.

#### The CX neural model

The CX model circuitry and input signals are described in Extended data figure 2 (a-d), and the different parameters used to obtain the output (motor command) are described in Extended data figure 1. All the modelling has been achieved with Matlab, and can be provided upon request.

---

**Extended data figure 1.**
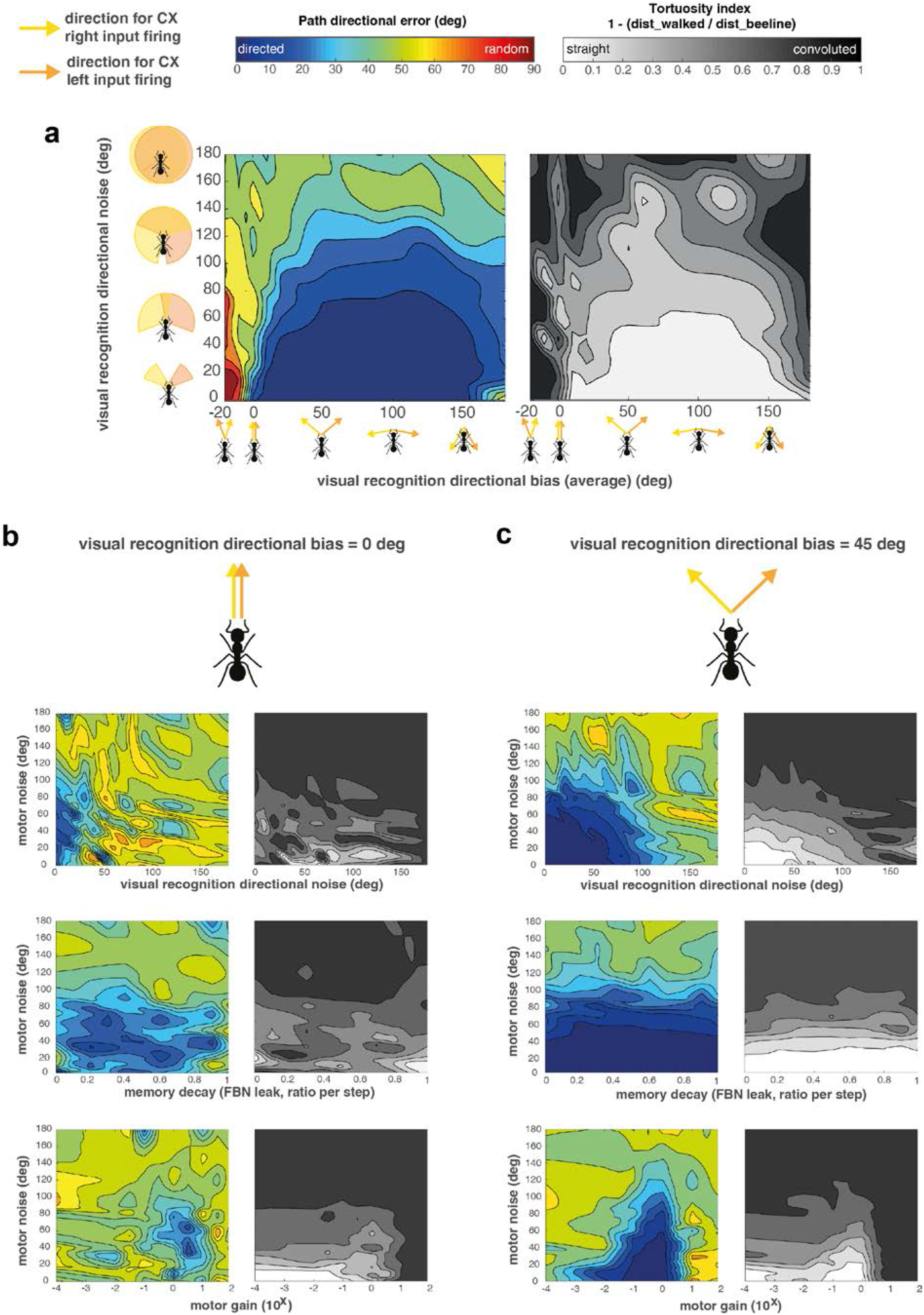
Parameter exploration of the Central Complex model (see fig 3). **a**. This shows a parameter exploration for the CX model presented in **Fig. 1** (see **extended fig. 2** for details of the circuitry). – Path directional error (absolute angular error between start-to-arrival and start-to-goal directions) and path tortuosity (index = *1 − (beeline_distance/distance_walked)*) after 200 steps are shown according to various parameter ranges. For each point on the map, all the other parameters are chosen to maximise for lowest path directionality error. **a**. Same as **Fig. 1d**, except that for each point of the map, the other parameters are chosen to maximise for lowest path directionality error instead of being fixed at an average range. Note that in **Fig. 1d**, visual familiarity direction bias < 0 typically results in routes leading to the opposite direction (i.e., path directional error close to 180°, **see Fig. 1**). Here, maximising for lowest path directional error did not result in goal-oriented path, but selected parameters yielding very high tortuosity, thus indicated that no parameter regime can yield straight, directed route when visual familiarity bias is < 0. Note that straight, goal-oriented paths emerge as long as the visual familiarity direction bias is > 0, that is, if the left hemisphere inputs correlate with moments when the nest is on the left, and vice versa. **b**. Visual familiarity directional bias is fixed at a value of 0°, meaning that both CX inputs respond maximally when the agent is facing the goal direction. Note that in this condition, regions of low path directional errors (blue) and region of low path tortuosity (white) do not overlap. This means that one cannot obtain straight, goal-directed paths if left and right CX inputs respond when the nest is located in front. **c**. Visual familiarity directional bias is fixed at a value of +45°, meaning that left and right CX inputs respond maximally when the agent is oriented 45° to the right or left from the goal direction, respectively. Note that regions with low path directional errors (blue) and regions of low path tortuosity (white) overlap well, showing a very large range of parameters for which we can obtain straight, goal-directed paths. We found the robustness to parameters remarkable: the model copes with motor noise up to 80°, visual familiarity direction noise up to 90°, is insensitive to its vector-memory decay and operates across several orders of magnitude for the gain.

## Parameters’ description

***Visual familiarity directional bias*** Indicates the absolute angle away from the goal at which visual familiarity signals (i.e., the CX inputs) are highest, assuming 0° indicates the correct goal direction. 0° indicates that both left and right inputs fire when the nest direction is aligned with the current body orientation. Inversely, 180° indicates that left and right input fire when the nest is right behind. Positive values (between 0° and 180°) indicate that the left and right inputs fire when the nest direction is on the left and right hand side respectively (the extent of the angular bias is given by the value). Negative values (between 0° and −180°) indicate a reversal, so that left and right input fire when the nest direction is on the right and left hand side respectively. ***Visual familiarity directional noise:*** Represents the extent of a systematic deviation from the visual familiarity directional bias angle. It is implemented by shifting the input curve response (horizontal arrows in **Fig. 1b**) around its mean (given by the ‘directional bias’) at each time step by random values drawn from a normal distribution with standard deviation given by ‘directional noise’. It can be seen as representing a directional noise when storing visual memories. High directional noise means that the input signal will occasionally respond strongest when oriented in the other direction than indicated by the visual familiarity directional bias. Robustness to visual familiarity directional noise indicates that the orientation of the body does not need to be precisely controlled during memory acquisition. ***Motor noise***: at each time step, a directional ‘noise angle’ is drawn randomly from a Gaussian distribution of *± SD = motor noise*, and added to the agent’s current direction. ***Memory decay***: proportion of Fan-shaped Body Neurons (FBN, see extended fig 2 for details) activity lost at each time step: For each FBN: Activity_(t+1)_ = Activity_(t)_ × (1 – *memory decay*). This corresponds to the speed at which the memory of the vector representation in the FBN decays. A memory decay = 1 means that the vector representation in the FBN is used only for the current time step and entirely overridden by the next inputs. A memory decay = 0 means that the vectors representation acts as a perfect accumulator across the whole paths (as in PI), which is probably unrealistic. ***Motor gain***: Sets the gain to convert the motor neuron signals (see extended fig 2 for details) into an actual turn amplitude (*turn amplitude = turning neuron signal × gain*). Note that here, the motor gain is presented across orders of magnitude. One order of magnitude higher means that the agent will be one order of magnitude more sensitive to the turning signal.

**Extended data figure 2.**
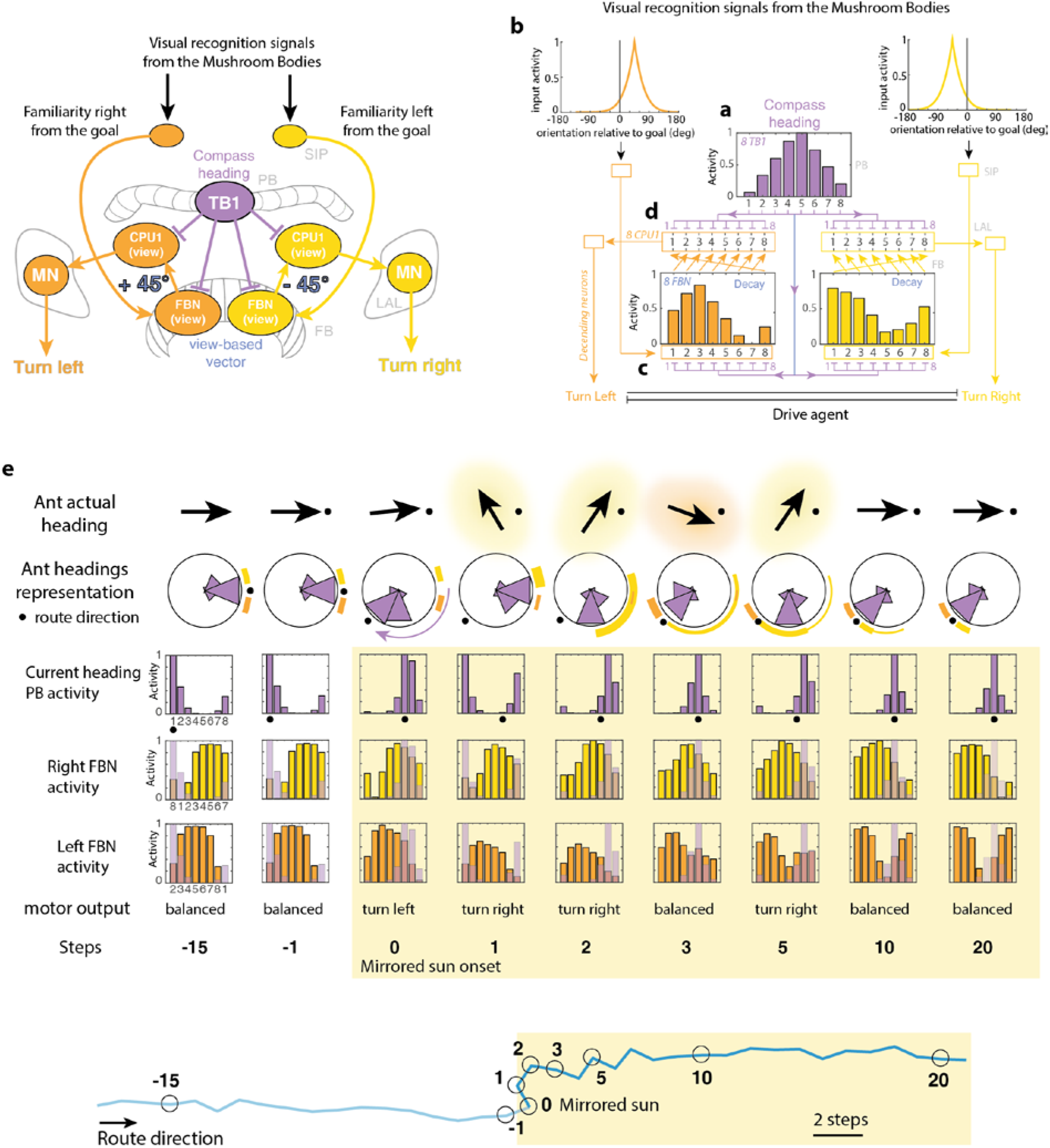
Details of the CX model’s circuitry. **a-d**. General scheme of the CX model as presented in figure 1 (left panel) and the corresponding detailed circuitry (right panel). This model exploits the same circuit as the CX model used for PI ^12,14^, except that FB input indicate visual familiarity rather than speed of movement. **a**. Current heading direction is modelled in the Protocerebral Bridge (PB) as a bump of activity across 8 neurons forming a ring-attractor (purple), as observed in insects ^14^. Each neuron responds maximally for a preferred compass direction, 45° apart from the neighbour neurons (neuron 1 and 8 are functionally neighbours, closing the ring structure). Change in the agent’s current compass orientation results in a shift of the bump of activity across the 8 neurons (we did not model how this is achieved from sensory cues, see ^9,10,61^ for studies dedicated on this. **b**. Visual familiarity signals fire according to the agent orientation relative to the goal direction. Here the input curve indicates that right and left signals fire maximally when the agent is oriented 50° (in average) left and right from its goal respectively (but see Fig. 1 and Extended fig. 1 for variation of these parameters: ‘directional bias’ and ‘directional noise’). **c**. These lateralised input signals excite two dedicated sets of FBN. These FBNs are simultaneously inhibited by the current heading representation (purple), resulting in two negative imprints of the current heading activity across the FBNs, which can be viewed as two ‘view-based vectors’. FBNs show some sustained activity so that, across time, successive imprints are superimposed, thus updating the ‘view-based-vectors’ (as for Path integration, except that this sustained activity is not crucial). The sustainability of such a ‘view-based vector’ depends on the FBN activity’s decaying rate, which can be varied in our model and has little incidence on the agent’s success (Extended figure 1, parameter decay). **d**. Motor control is achieved using the same circuitry as for Path integration ^12^. On each brain hemisphere, neurons (called CPU1 in some species), compare the current compass heading (purple) with their version of the FBN ‘view-based-vector’. Crucially, both FBN representations are neurally shifted by 1 neuron (as if rotating the view-based-vector by 45° clockwise or counter-clockwise depending on the hemisphere), resulting in an overall activity in the CPU1 (sum of the 8 CPU1) indicating whether the view-based-vector points rather on the left-(higher resulting activity in the left hemisphere) or right-hand side (higher resulting activity in the right hemisphere). The CPU1 neurons sum their activity on descending motor neurons (MN), which difference in activity across hemispheres triggers a left or right turn of various amplitude, given a ‘motor gain’ that can be varied to make the agent more or less reactive (Extended figure 1 for detailed parameter description). Numbers on the left indicate neurons numbers. Letters on the right indicate brain areas (SIP: Superior Intermediate Protocerebrum, PB: Protocerebral Bridge, FB: Fan-shaped Body, LAL: Lateral Accessory Lobe). **e**. Same as Fig. 4c, with added details of the PB (purple) and right and left FB (yellow and orange) neural activity. Note that the FBNs order has been shifted (2,3,4,5,6,7,8,1 and 8,1,2,3,4,5,6,7) and inhibition exerted by the PB is represented (overlaid transparent purple, 1,2,3,4,5,6,7,8) as happens in the left and right CPU1 neuron (**d**). This way, the strength of the motor signal for turning right and left– which correspond to the sum of non-inhibited right and left CPU1 activity – can be inferred by looking at the area covered by non-occluded yellow and orange FBN columns respectively. With manipulation such as rotating the current compass information, it becomes apparent that motor decision results from complex dynamics between two main factors: 1- how strong are the left and right visual input signal updating the view-based-vectors representation (represented by orange and yellow glow around the actual ant heading arrows), which depend on whether the agent is oriented left or right from its goal and 2- how well the current heading representation (PB) matches the rotated left and right shifted FB view-based-vector current representations.

